# Graspot: A graph attention network for spatial transcriptomics data integration with optimal transport

**DOI:** 10.1101/2024.02.01.578505

**Authors:** Zizhan Gao, Kai Cao, Lin Wan

**Affiliations:** Academy of Mathematics and Systems Science, Chinese Academy of Sciences, Beijing, China; Eric and Wendy Schmidt Center, Broad Institute of MIT and Harvard,Cambridge, USA

**Keywords:** spatial transcriptomics, data integration, optimal transport, graph attention neural network

## Abstract

Spatial transcriptomics (ST) technologies enable the measurement of mRNA expression while simultaneously capturing spot locations. By integrating ST data, the 3D structure of a tissue can be reconstructed, yielding a comprehensive understanding of the tissue’s intricacies. Nevertheless, a computational challenge persists: how to remove batch effects while preserving genuine biological structure variations across ST data. To address this, we introduce Graspot, a **gr**aph **a**ttention network designed for **sp**atial transcriptomics data integration with unbalanced **o**ptimal **t**ransport. Graspot adeptly harnesses both gene expression and spatial information to align common structures across multiple ST datasets. It embeds multiple ST datasets into a unified latent space, facilitating the partial alignment of spots from different slices. Demonstrating superior performance compared to existing methods on four real spatial transcriptomics datasets, Graspot excels in ST data integration, including tasks that require partial alignment. In particular, Graspot unveils subtle tumor microenvironment structures of breast cancer, and accurately aligns the spatio-temporal transcriptomics data to reconstruct human heart developmental processes. The code for Graspot is available at https://github.com/zhan009/Graspot.

## Introduction

Recently, spatial transcriptomics (ST) technologies have enabled the simultaneous measurement of mRNA expression while retaining spatial information within a tissue slice. These techniques have been extensively applied to investigate both healthy regions [3] and diseased tissues, including cancerous tissues [12, 17]. The integration of multiple slices provides a comprehensive understanding of complex biological processes in entire tissues, fostering innovative approaches to downstream tasks such as analyzing spatial expression across slices and exploring 3D cell-cell communications. However, it remains challenging to jointly analyze multiple ST adjacent slices due to batch effect and biological structure variations across slices. Several methods have been used for alignment and integration of multiple ST slices. For example, PASTE [19], which is based on fused Gromov–Wasserstein optimal transport, can align slices in a global manner. However, PASTE assumes that slices have global similarity and fails to consider the biological structure variations across slices. To solve the problem, PASTE2 [14] extends the single-cell data integration method Pamona [7] and introduces a partial fused Gromov-Wasserstein method to incorporate spatial information. Meanwhile, both PASTE and PASTE2 work on original space or linear embedding, and cannot provide a common integrated space across ST slices.

Moreover, nonlinear embedding based methods are developed, such as GPSA [13] and STAligner [20]. GPSA [13] integrates multiple ST slices into a common coordinate system using deep Gaussian processes, while STAligner [20] embeds gene expression and spatial neighbor network of spots with a graph attention network and aligns slices using the mutual nearest neighbor (MNN) originally developed for single-cell data integration. However, none of them are capable of handling the partial alignment of slices or providing the probabilistic mapping of spots in adjacent slides for downstream analysis, as achieved by PASTE and PASTE2. There are also some generic methods, such as Tangram [5] and uniPort [6], are designed to integrate and align single-cell and ST data. Tangram [5] is a deep learning method that aligns single-cell data onto ST data, while uniPort [6] is a unified single-cell data integration framework that combines a coupled variational autoencoder and mini-batch unbalanced optimal transport (UOT). The UOT module allows uniPort more suitable for heterogeneous data integration. However, both Tangram and uniPort fail to utilize the spatial information provided by ST data.

In this work, we present Graspot, a graph attention network with unbalanced optimal transport that aligns and integrates multiple spatial transcriptomics data. Graspot extends uniPort by leveraging both graph attention network (GAT) and UOT with the aim to efficiently utilize both gene expression and spatial information provided by ST data for the integration. Graspot inherits the graph attention network architecture of STAGATE [10] which nonlinearly embeds both gene expression and spatial information of each slice into a common latent space. The UOT module of Graspot enables it to reconstruct the probabilistic mapping of spots in the latent space across slices [16]. Based on the partial probabilistic mapping, Graspot not only efficiently removes the batch effect, but also retains the biological structure variations across slices. We iteratively optimize the graph attention network and UOT distance to generate both the integrated embedding and probabilistic alignment.

We verified Graspot on human dorsolateral prefrontal cortex (DLPFC) ST data, anterior sagittal mouse brain ST data, HER2 breast cancer ST data and ST datas of Human heart development. Our results indicate that Graspot attains highest alignment accuracy compared to competing methods both on global and partial alignment. Graspot can also uncover subtle structures within the tumor microenvironment from multiple slices. In addition, Graspot accurately aligns spatio-temporal ST slices to reveal the human heart developmental processes.

## Materials and methods

### Overview of Graspot

Graspot is a graph attention network for spatial transcriptomics data integration with unbalanced optimal transport. It is comprised of two modules: the graph attention network (GAT) module and the unbalanced optimal transport (UOT) module.

The main inputs of Graspot are multiple spatial transcriptomics slices which contain both gene expression profiles and two-dimensional spot coordinates. There are two main outputs of Graspot: (i) a probabilistic alignment of spots across multiple spatial transcriptomics slices, and (ii) a common low-dimensional space that recovers and aligns multiple spatial transcriptomics slices.

### Mathematical formulation of Graspot

Suppose that two ST slices (*x*_1*i*_, *p*_1*i*_), *i* = 1, 2, …*n*_1_ and (*x*_2*j*_, *p*_2*j*_), *j* = 1, 2, …*n*_2_ are inputs of Graspot, where *x*_1*i*_ ∈ ℝ^*g*^ is the normalized gene expression of spot *i, x*_2*j*_ ∈ ℝ^*g*^ is the normalized gene expression of spot *j* and *p*_1*i*_(*p*_2*j*_) ∈ ℝ^2^ is the coordinate of spot *i*(*j*). *g* represents for the number of highly variable common genes. Graspot first builds an undirected spatial neighbor network and computes an adjacency matrix *A*_*ij*_ based on spot locations in respective slices. Here, *A*_*ij*_ = 1 if and only if the Euclidean distance between spot *i* and spot *j* is less than a predefined radius *r*. Afterwards, Graspot inputs two ST slices through a graph attention network (GAT) framework and learns its *k*-dimensional embeddings. The latent vector *z* generated by the graph attention auto-encoder which achieves nonlinear dimensionality reduction while collectively aggregating information from neighbors of spots.

The neural network module in Graspot consists of three parts: an encoder, a decoder and the graph attention layers. The inputs of encoder are initialing as *h*^(0)^ = *x* = (*x*_1_, *x*_2_)^*T*^, *x*_*i*_ ∈ ℝ^*g*^, *i* = 1, 2, …, *n*_1_, *n*_1_ + 1, …, *N* where *N* = *n*_1_ + *n*_2_. Encoder in Graspot generates the embedding of spot *i* in *t*-th layer as follows:

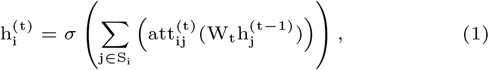

where σ is the nonlinear activation function, *S*_*i*_ is the neighbor set of spot *i* according to spatial neighbor network, and *W*_*t*_ is weight matrix for training. Besides, Graspot employs a self-attention mechanism, which is an added single-layer neural network in *t*-th layer in the encoder, to adaptively learn information from neighbors. The edge weight between spot *i* and spot *j* from same slice is described as follows:

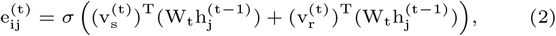

Where 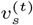 and 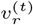 are trainable weight vectors, 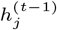 is the represent of spot *j* in *t* − 1 layer, and *att*_*ij*_ is normalized form of *e*_*ij*_ via a softmax function as follows:

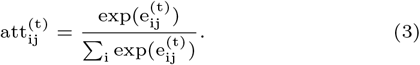

After obtaining the latent representations from the encoder, Graspot applies a decoder to reverse the final spot embedding back to the original normalized gene expression. The last layer of the decoder is formulated as follows:

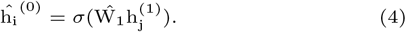

It has the same number of layers as encoder and we set 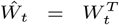 and 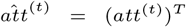 respectively to avoid overfitting. One of the objectives minimized by Graspot is the reconstruction loss of normalized expressions as follows:

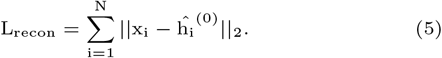

To better integrate heterogeneous ST slices in a common low-dimensional space, we further design an alignment term for different ST slices using Unbalanced Optimal Transport. Suppose we have two latent vectors 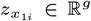 and 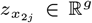 generated by encoder *ψ*, respectively. Due to the favorable properties of the common latent space that recover intrinsic structures of ST slice, the UOT cost between spot *i* and spot *j* in different slices is defined as Euclidean distance of latent vectors 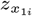 and 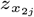 as follows:

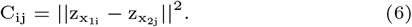

We then compute the following unbalanced entropic optimal transport plan [4]:

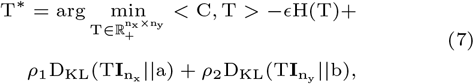

where < *C, T* >=Σ_*i,j*_ *C*_*ij*_ *T*_*ij*_, entropy regularization term *H*(*T*) = − *Σ* _*i,j*_(log *T*_*ij*_ − 1), and 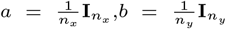. *ϵ* is an entropy parameter. *ρ*1 and *ρ*2 are two regularization parameters. As mentioned above, multiple ST slices may only partially overlap, and as a result, there may be a portion of spots in both slices that are seldom aligned. It is worth noting that the UOT module in Graspot allows only a fraction of the total mass to be transported between two distributions. Eq. (7) combines entropic regularization [8] with a more general formulation for unbalanced optimal transport. Unbalanced entropic optimal transport allows to slightly diverge from initial mass during transportation and can be optimized using variations of the Sinkhorn algorithm [11]. This is a strictly convex optimization problem and can be solved via an efficient inexact proximal point method [18] as:

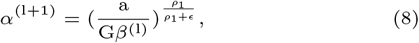

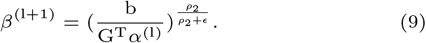

The initial 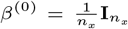 and 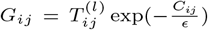, and the UOT plan 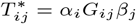. The second objective of our method is defined as the UOT loss:

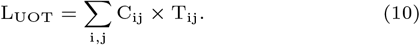

Therefore, the total objective minimized by Graspot is formulated as:

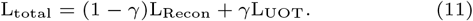

Here, *γ* is the trade-off of reconstruction loss and unbalanced optimal transport loss. We set *γ* = 0.2 in the following experiments.

#### Algorithm 1

Graspot: a graph attention network for spatial transcriptomics data integration with optimal transport.

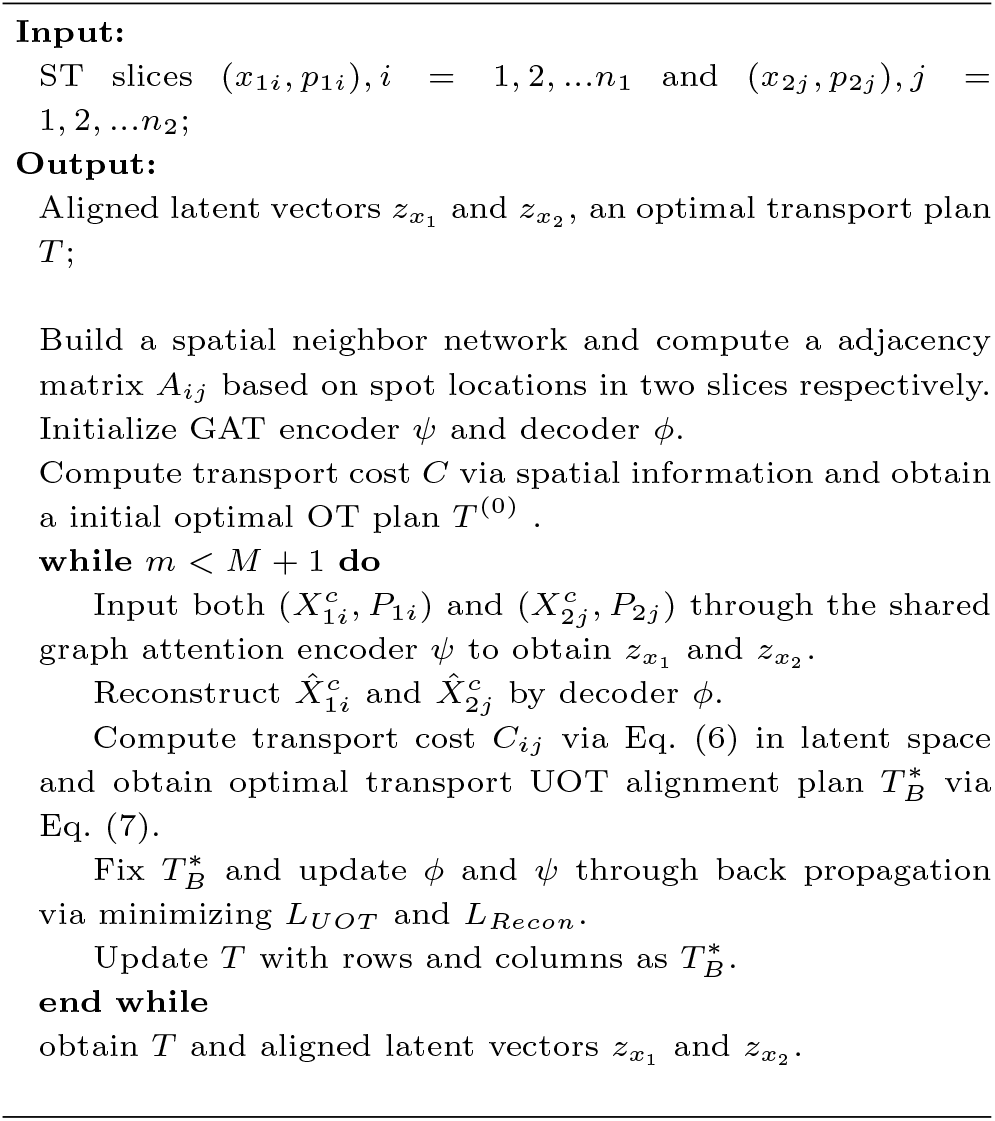

We perform UOT and graph attention auto-encoder training in an iterative manner to generate a probabilistic optimal transport plan and a common aligned low-dimensional space.

Finally, the optimal transport plan *T* produced as the output represents the probabilistic correspondence between spots. The encoder’s output is a commonly aligned low-dimensional space for multiple ST slices. We first initialize *T* ^(0)^ as the optimal transport plan of the optimal transport problem based on the spatial information as in PASTE [19]. Once *T* is obtained, Graspot provides users an option to initialize *T* ^(0)^ in the optimization of Eq. (7) to enhance the participation of spatial information in alignment module. Details of Graspot algorithm are shown in Algorithm 1.

## Methods evaluation metrics

We evaluate our methods both from the spots mapping *T* for alignment and nonlinear integrated embedding *z*. To evaluate spots mapping *T* for alignment, we employ the metrics of Alignment accuracy and Label Transfer ARI as follows.

### Alignment accuracy

We assume that spot *i* is in slice A and spot *j* is in slice B, and Slice A has *n*_1_ spots and slice B has *n*_2_ spots. The Alignment accuracy evaluation metric is defined as follows:

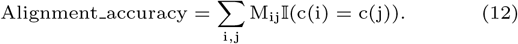

When *n*_1_ < *n*_2_, we compute *M*_*ij*_ by row as follows:

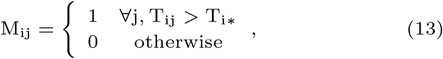

where *T*_*i*\*_ is the *i*-th row in mapping *T* and *c*(*i*)*/c*(*j*) indicates the annotated or clustered regions which spot *i*/spot *j* belongs to.

### Label Transfer ARI

Given *M*_*ij*_ which indicates the maximum probability matching, we compute the one-to-one spot correspondence π : *n*_1_ → *n*_2_. The Label Transfer ARI metric is defined as ARI score of cell-type clusters in sliceA and reorder cell-type clusters in sliceB derived from spot correspondence π(*i*), *i* = 1, …, *n*_1_. The Adjusted Rand Index (ARI) is utilized to assess the similarity between two different cell-type clusters. The ARI value ranges from 0 to 1, with 0 for random labeling and 1 for perfect matching.

To evaluate nonlinear integrated embedding *z*, we employ the metrics of two categories: (1) removal of batch effects and (2) conservation of biological structure variations. Here the former category includes Batch Entropy score and Batch Average Silhouette Width (Batch ASW) and the latter category includes Silhouette (Cell-type ASW) score.

### Batch Entropy

Batch entropy score is derived from uniPort [6] which evaluates the sum of regional mixing entropies at the location of randomly chosen spots from different slices. It can be calculated as:

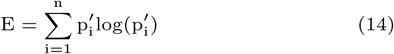

Where

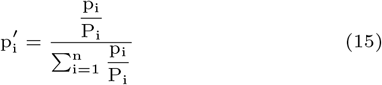

*p*_*i*_ is the proportion of spot numbers in each batch to the total spot numbers, and *P*_*i*_ is the proportion of spots from batch *i* in a given region. A high Batch entropy score indicates spots from various slices mixing well.

### Batch ASW

Batch Average Silhouette Width(Batch ASW) refers to modified silhouette width to measure batch mixing. The Alignment accuracy evaluation metric is defined as follows:

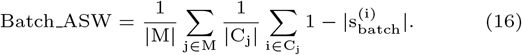

where *M* is the set of unique spot labels, *C*_*j*_ is the set of spots with the spot label *j* and *C*_*j*_ denotes the number of spots in that set. |*s*_*batch*_(*i*)| is the silhouette width on batch labels for the *i*-th spot. Higher Batch ASW indicates ideal mixing.

### Silhouette

Silhouette (Cell-type ASW) is used to determine the separation of spot clusters. It is defined as follows:

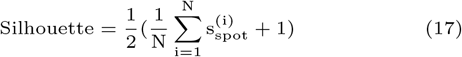

where *N* presents the total number of spots and 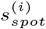 is the silhouette width on spot cluster labels for the *i*-th spot. Higher score of Silhouette represents better biology conservation of the slice integration.

## Results

### Graspot outperforms the state-of-the-art methods on global alignment with highest accuracies

We applied Graspot to analyze 10X Genomics Visium ST data from human dorsolateral prefrontal cortex (DLPFC). This dataset consists of 12 slices of DLPFC tissue from three samples, denoted as I, II and III. Each sample consists of four slices labeled from A to D along the *z*-axis. Within each sample, the first adjacent pair AB and the last adjacent pair CD are 10 *um* apart, whereas the middle pair BC of slices is located 300 *um* apart [15]. The slices were manually annotated into six neocortical layers plus a white matter by [15]. We used the annotation for method evaluations.

We applied Graspot to compute a pairwise slice alignment of each pair of consecutive slices from the same sample. We compared the pairwise alignments obtained by Graspot with competing methods: PASTE [19], STAligner[10] and Tangram[5]. The visualization of parallel spot matching results in Sample III using Graspot is shown in Fig. 2(a).

**Fig. 1.**
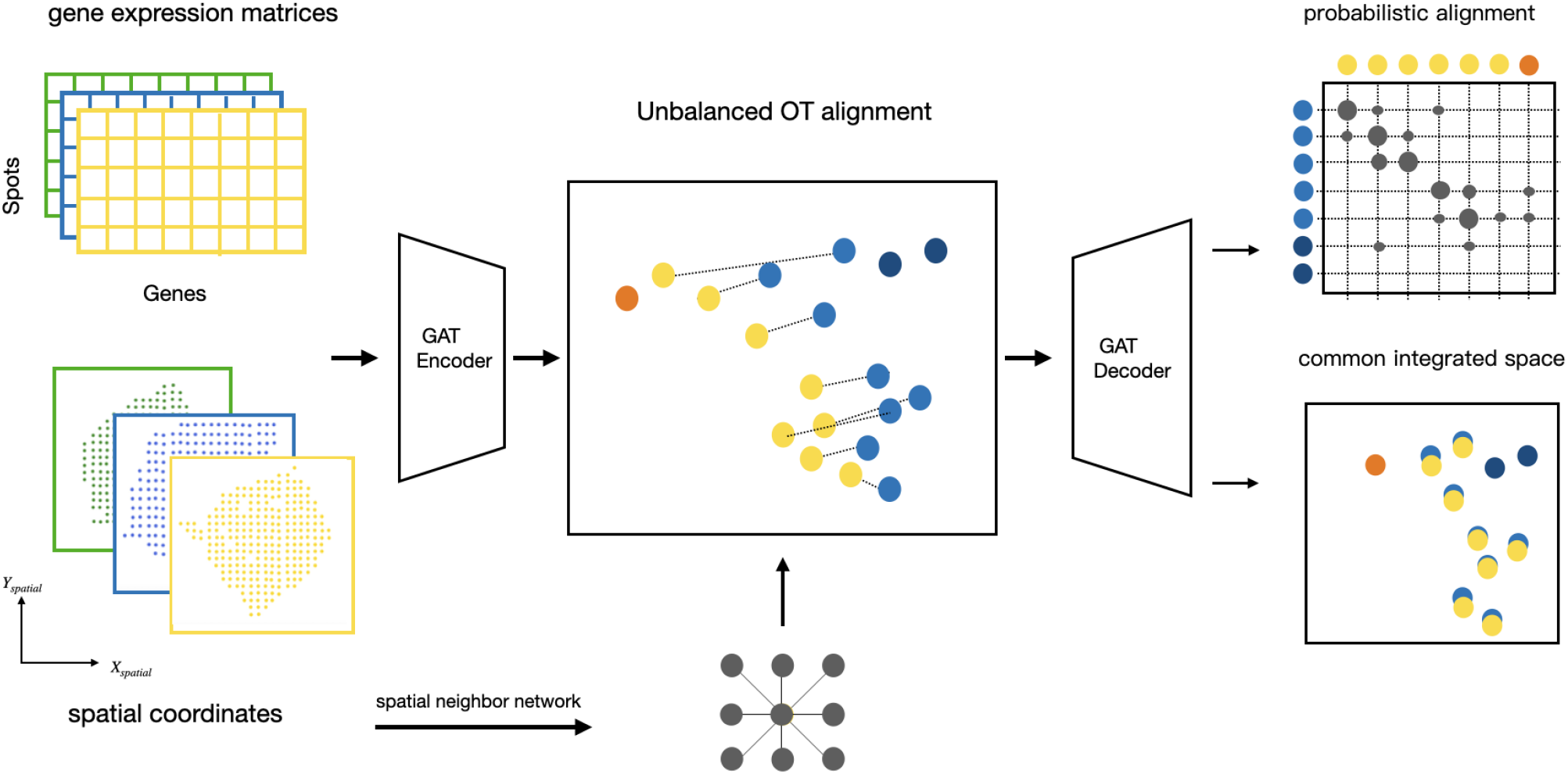
Overview of Graspot algorithm. Graspot is composed by two modules: graph attention network (GAT) module and UOT alignment module. The inputs of Graspot are multiple spatial transcriptomics slices with gene expression matrices and spatial coordinates. The outputs of Graspot are probabilistic alignment *T* and common integrated space aligns intrinsic structures of multiple ST slices.

**Fig. 2.**
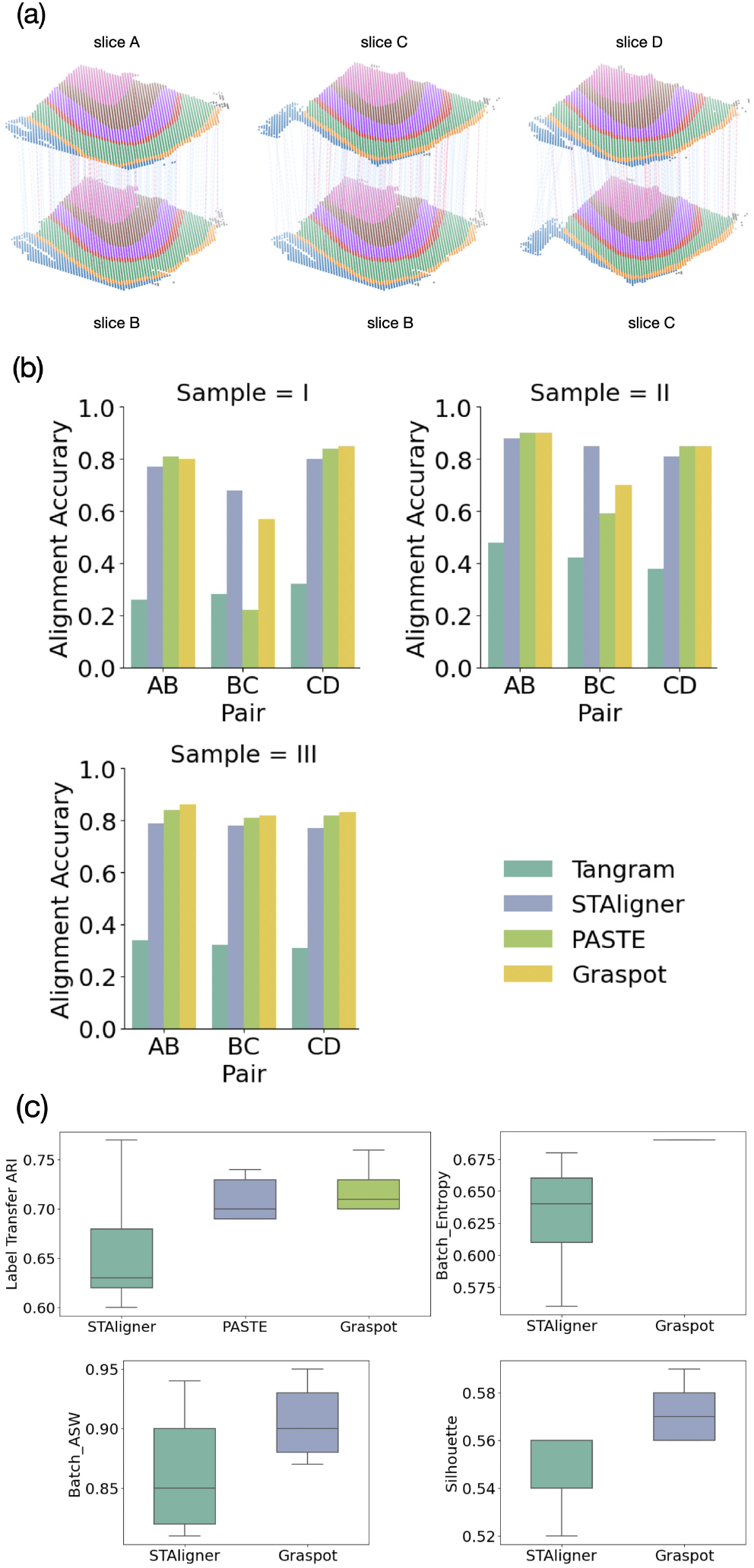
(a) Spot matching results in Sample III using Graspot. Blue lines indicate correct matches, whereas the red lines denote incorrect matches. (b) Accuracy of pairwise alignment of consecutive DLPFC slices (labeled AB, BC, and CD) for Tangram, STAligner, PASTE and Graspot, which is computed based on published annotations of spots. (c) Label Transfer ARI, Batch entropy, Batch ASW and Silhouette scores of pairwise slices alignment.

We evaluate Graspot both from the probabilistic spots mapping *T* for alignment and nonlinear integrated embedding. First, we measured the quality of spots matching by calculating the alignment accuracy that the fraction of matched spots belonging to the same annotated layer across slices and Label Transfer ARI which indicates the overlap of annotated layers and correspondent layers. Since STAligner can not provide the probabilistic alignment across slices, we performed matching strategy in its embedding space for the following evaluation. Second, we employed a series of scores including Batch entropy, Batch ASW and Silhouette to assess the performance of Graspot in embedding space. Due to the lack of a common integrated space in PASTE and Tangram, we only compared our method with STAligner.

We compared the other three algorithms mentioned above with Graspot and the results are shown in the Fig. 2(b,c). Graspot achieved the highest alignment accuracy in 6 of 9 pairwise alignment which shows the efficiency of our method in the alignment tasks in ST of human DLPFC data. As Fig. 2(b) shows, Graspot obtained a higher accuracy mostly above 0.8 in the close pairs (AB and CD) but lower accuracy in the far pairs (BC). Tangram algorithm achieved lowest accuracy from 0.26 to 0.48 on all slice pairs. STAligner algorithm achieved lower accuracy from 0.77 to 0.88 on all slice pairs, outperforming Graspot only on one middle BC slice pair in Sample I and II.

Graspot achieved higher Label Transfer ARI scores with less variation than other methods in 9 pairwise alignment tasks. The mean Label Transfer ARI score using Graspot is 0.67, surpassing other compared methods. Fig. 2(c) visualizes the results of Batch entropy, Batch ASW and Silhouette score. Graspot is more competitive than STAligner both on removing batch effects and conserving biological structure variations in latent embeddings.

These results demonstrate that Graspot, which combines transcriptional and spatial information to align ST data, frequently outperforms algorithms that use only transcriptional information. Additionally, our method, which iteratively learns the embedding and mapping using unbalanced optimal transport, exhibits better performance compared to the fused Gromov–Wasserstein optimal transport alignment strategy employed in PASTE and spot triplet learning alignment strategy in STAligner. Fig. 2 shows the superiority in evaluation of pairwise alignment of consecutive DLPFC slices (labeled AB, BC, and CD) for Tangram, STAligner, PASTE and Graspot. It is evident that Graspot improves alignment performance between adjacent slices. For instance, this enhancement is noticeable for all pairs in Sample III and for the BC pairs in both Sample I and Sample II.

### Graspot outperforms the state-of-the-art methods on partial alignment with highest accuracies

We applied Graspot to analyze 10x Genomics Visium ST data from human DLPFC tissues from three samples as the former experiment. To evaluate our method’s ability to handle unbalanced datasets using the UOT module, we tested Graspot on vertical partial subslices of human DLPFC tissues. The results are presented below. Following the strategy of matching to the closest in [9], we set *ρ*_1_ → ∞ and *ρ*_2_ = 0.01 for partial alignment. The slices that we selected are derived from four DLPFC slices in Sample III. For each slice, we created two partially overlapping subslices in the following manner: selecting two vertical lines on a slice to generate two subslices. The subslice on the left of line1 and the subslice on the right of line2 each encompass 70% of the total spots, as illustrated in Fig. 3(a).

**Fig. 3.**
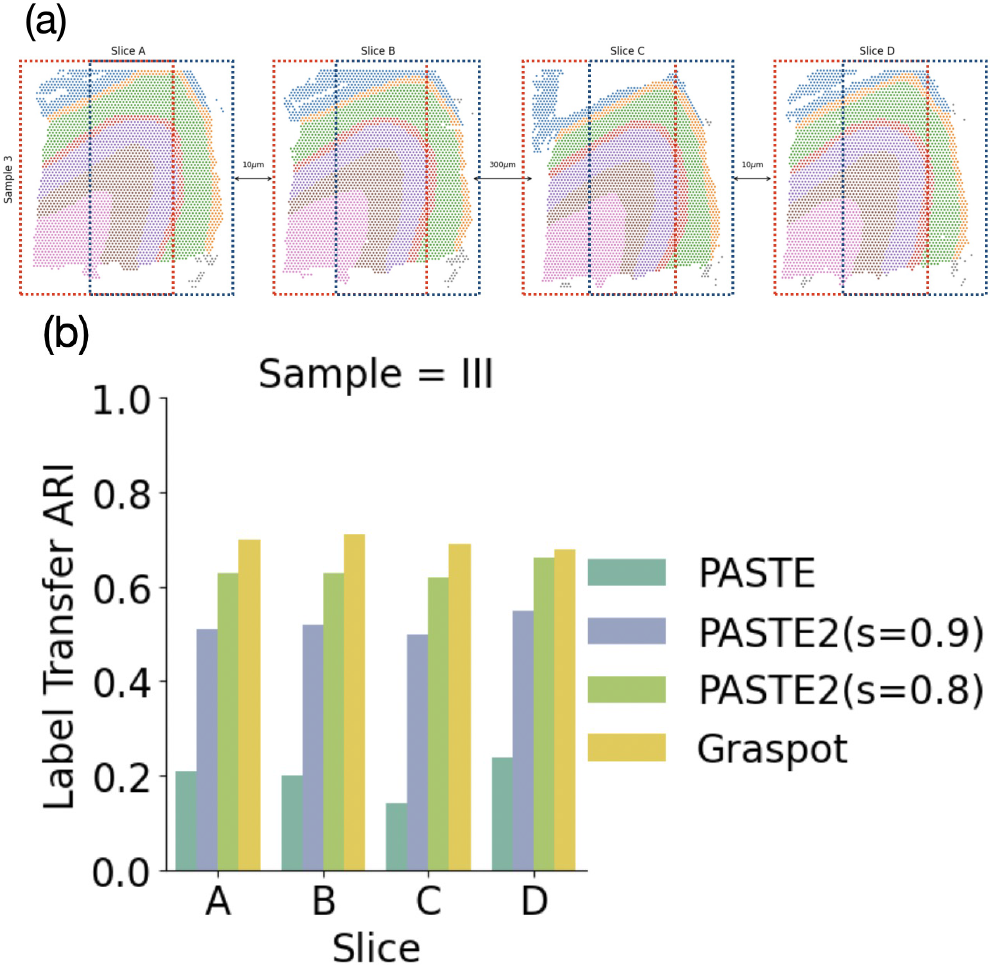
**(a) Two vertical subslices, one on the left and the other on the right, exhibit a 70% overlap of spots across four slices from Sample III. (b) Label transfer ARI scores of pairwise alignment of subslices in (a) for PASTE, PASTE2(s=0.8), PASTE2(s=0.9) and Graspot.**

Firstly, we applied our method to find alignments within the two created subslices for each of the slices A, B, C, and D in Sample III, separately. We follow PASTE2 [14] to use the Label Transfer ARI to measure the accuracy of partial alignment: Ground Truth clustering in slice with fewer spots and reordered Ground Truth clustering induced from spot correspondence π(*i*), *i* = 1, …, *n*_1_ in another slice. We compared our results with the other two algorithms (Fig. 3(b)). Consequently, Graspot demonstrated superior performance with results ranging between 0.68 to 0.71. In contrast, PASTE exhibited lower alignment accuracy, with values ranging from 0.14 to 0.24, attributable to its inability to handle partially overlapping slices. PASTE2 is a method designed for partial alignment, allowing only a fraction *s* of the probability mass to be transported. Here we set *s* equal to 0.8, PASTE2 addressed a partial fused Gromov-Wasserstein optimal transport problem but achieved a lower alignment result, ranging from 0.62 to 0.66, compared to Graspot. PASTE2 yielded inferior outcomes with *s* = 0.9, varying from 0.50 to 0.55. This suggests that PASTE2, when set to a larger *s* value, intends to align the entirety of spots between slices.

### Graspot efficiently integrates multiple ST slices

We applied Graspot to integrate ST data from four slices of human DLPFC Sample III. Once trained, Graspot has the capability to iteratively integrate multiple slices into an aligned embedding within a common low-dimensional space. Using the graph attention neural network after the alignment of Slice B and Slice C in Sample III, we conducted an experiment to demonstrate that Graspot integrates multiple slices in an iterative manner. We tested Graspot on slices from Sample III. Firstly, we needed to train the neural network thoroughly within our model to align the pairwise slices (Slice B and Slice C) and then input the new slices from the same sample (Slice A and Slice D). As shown in Fig. 4(a), the newly inputted slices, including Slice A and Slice D, align well with the existing slice embeddings in the common low-dimensional space. Additionally, the embeddings display clear patterns of cluster results that are in accordance with manually annotated layers. We also integrated the 4 slices in a common space using STAligner for comparison (Fig. 4(b)). The embeddings produced by Graspot demonstrate superior alignment and distinct clustering when compared to STAligner, particularly in cases involving the integration of multiple spatial transcriptomics slices.

**Fig. 4.**
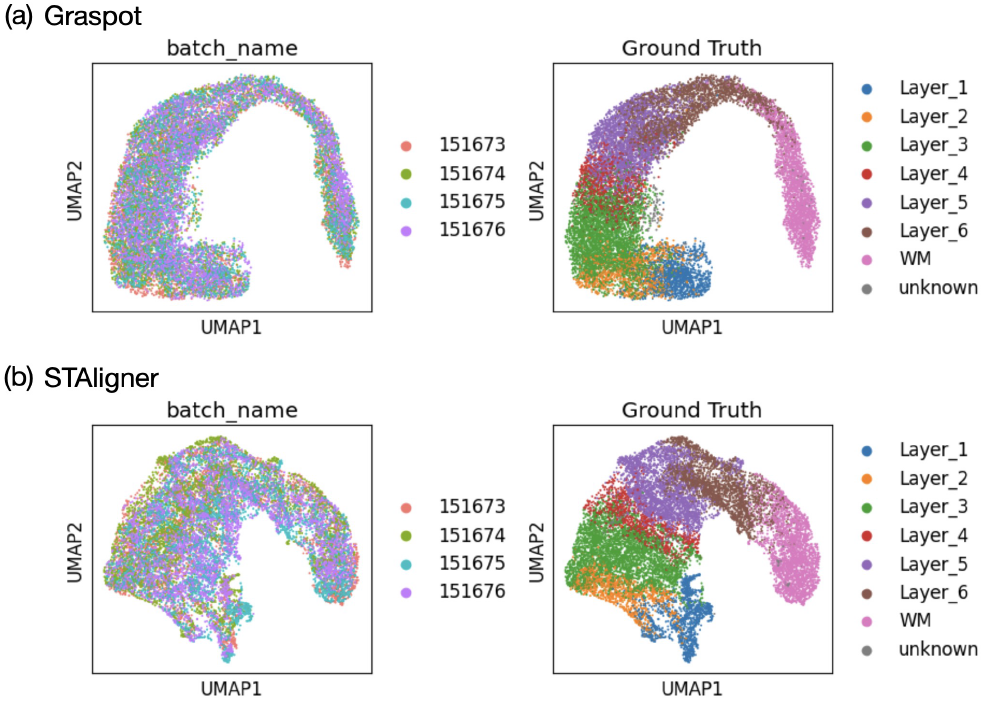
(a) Left: Embedding of four slices in Sample III using Graspot. Right: Clustering results of Graspot. (b) Left: Embedding of four slices in Sample III using STAligner. Right: Clustering results of STAligner.

We also applied Graspot to analyze three ST slices from HER2 breast cancer of patient G [1]. Our experiment demonstrated that Graspot is capable of effectively performing tasks related to alignment, integration, and identification of distinct spatial structures in biological data. The three ST slices from HER2 breast cancer of patient G are adjacent and part of slices are partial overlap, as depicted in Fig. 5(a). Following the alignment of the pairwise slices G1 and G2 using Graspot, the subsequent slice G3 aligns effectively with the existing latent embedding in the shared low-dimensional space. This process results in an integrated embedding, as depicted in Fig. 5(b). Afterwards, we employed the Louvain clustering algorithm on integrated spots within the common low-dimensional space. The clustering outcomes for the template slice G2 are displayed in Fig. 5(c). Graspot successfully identifies two regions of spots, aligning with the in situ cancer regions identified by pathological annotations, as illustrated in Fig. 6(d). These results affirm that the integration of multiple spatial transcriptomics slices using Graspot effectively preserves and unveils intricate structures within the tumor microenvironment.

**Fig. 5.**
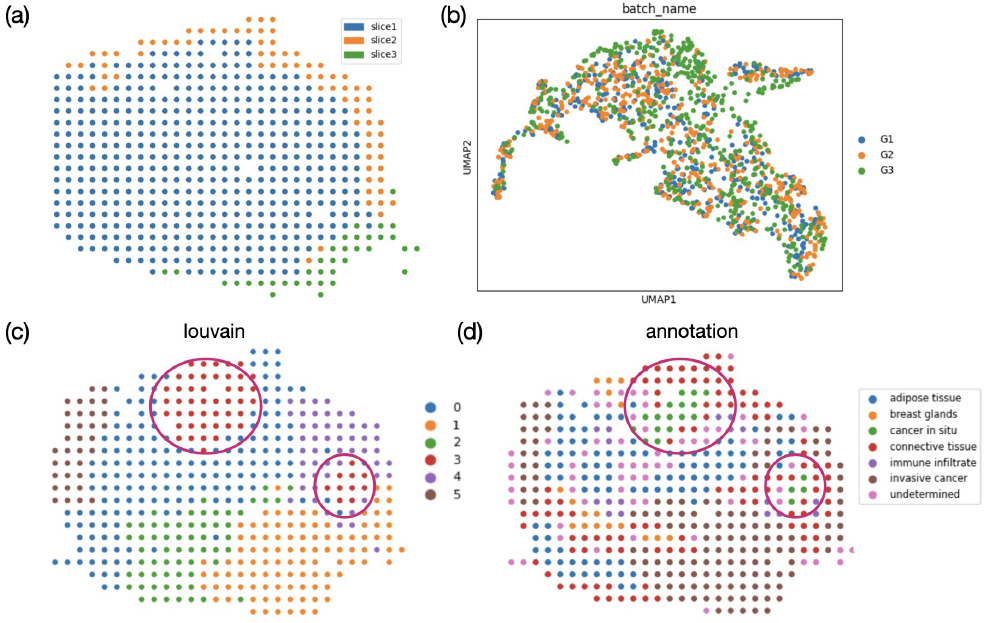
Graspot integrated multiple human breast cancer spatial transcriptomics data. (a) Three spatial transcriptomics slices from HER2 breast cancer patient G with sections of adjacent slices overlapping along the z-axis. (b) Integrated embedding of three ST slices in common space. (c) Clusters in template slice G2. (d) Pathological annotations of G2 slice.

**Fig. 6.**
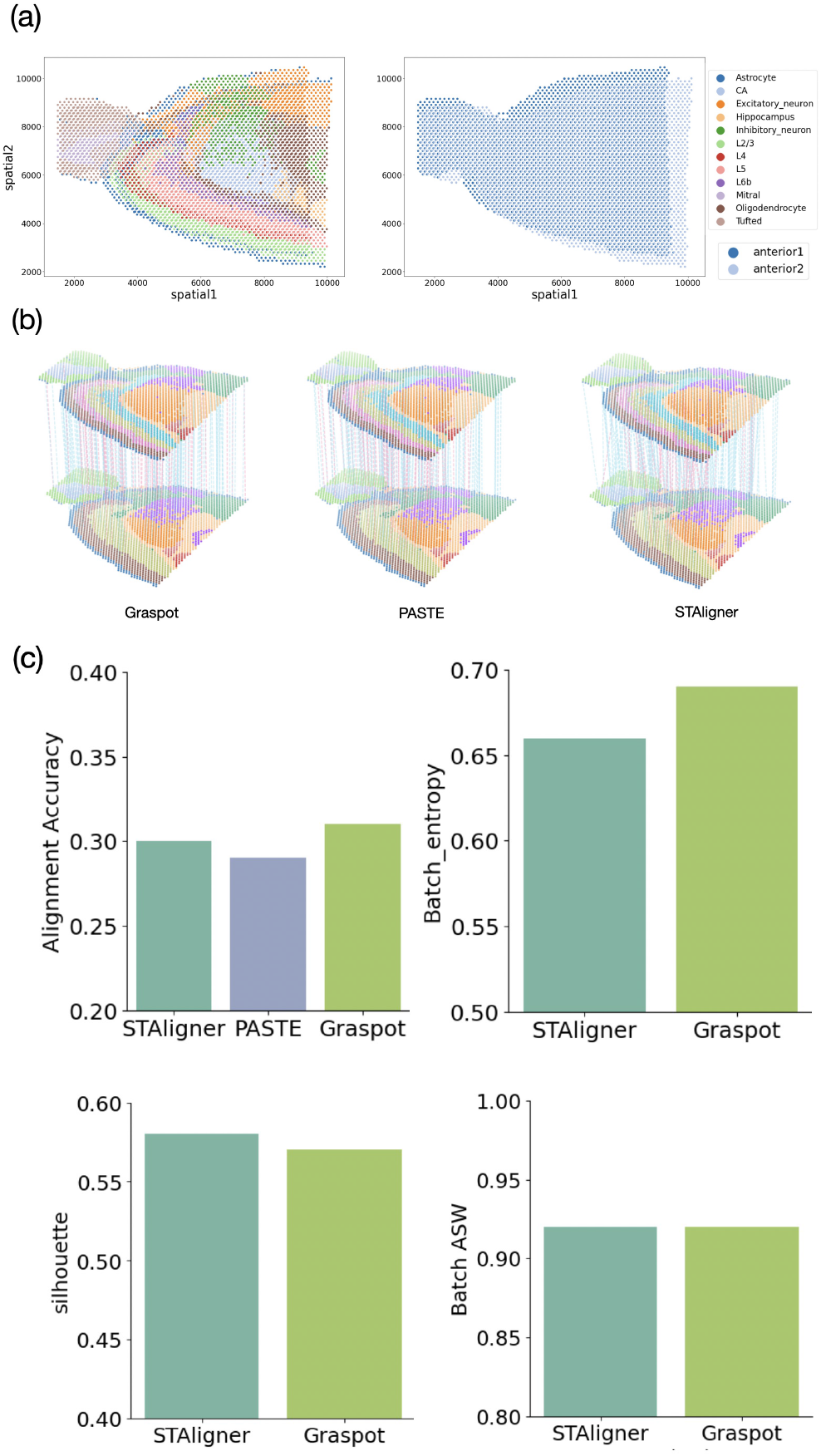
(a) A pair of replicate spatial transcriptomics slices from the anterior sagittal section of a mouse brain with complex structures, overlapping along the z-axis. (b) Spot matching results in Sample III using Graspot, PASTE and STAligner. Blue lines indicate correct matches, whereas red lines denote incorrect matches. (c) Label Transfer ARI, Batch entropy, Batch ASW and Silhouette scores of pairwise slices alignment.

### Graspot efficiently integrates ST slices from anterior sagittal mouse brain

We applied Graspot to analyze 10X Genomics Visium ST data from sagittal mouse brain with complex structures. Sagittal mouse brain ST slices consist of two pairs of replicates in both anterior and posterior positions separately. Figure 6(a) indicates the two replicates are most partially overlap with similar complex structures. Here we take one pair of sagittal mouse brain ST datas which labeled as “anterior 1” and “anterior 2” for our experiment. Fig.6(b) depictes the 3D landscape of spot matching results between two replicates of sagittal mouse brain data using Graspot, PASTE and STAligner, respectively.

We benchmarked the performances of methods using the ground truth of annotations which contain common hierarchical layers including “L5”, “L2/3”,”Astrocyte” and “CA”. We conducted alignment accuracy evaluation in probabilistic spots mapping *T* compared with PASTE and STAligner. Graspot achieved a highest score of 0.31, while STAligner achieved the second highest score of 0.30, and PASTE achieved a lowest score of 0.29 (upper-left panel of Fig. 6(c)).

To assess and demonstrate the integration performance within the common space, we used metrics such as Batch entropy, Batch ASW, and silhouette score for comparing two methods, namely STAligner and Graspot. Given that PASTE does not offer a common embedding space, it was excluded from these subsequent comparisons. We observed that Graspot obtain a higher Batch entropy score than that of STAligner (upper-right panel of Fig. 6(c)), while its silhouette score is slightly lower than STAligner (lower-left panel of Fig. 6(c)). Meanwhile, Graspot and STAligner have the same Batch ASW scores shown in lower-right panel of Fig. 6(c).

### Graspot partially aligns spatio-temporal ST datas of Human heart development

We applied Graspot to explore the changes of spatio-temporal distribution of different spot types during embryonic development using ST datas collected from human heart in early embryos, ranging from 4.5–5 post-conception weeks (PCW), 6.5 PCW to 9 PCW [2]. There are four, nine, and six tissue slices sampled along the dorsal-ventral axis from the 4.5–5PCW, 6.5PCW, and 9 PCW heart tissues, respectively (Fig.7(a)).

**Fig. 7.**
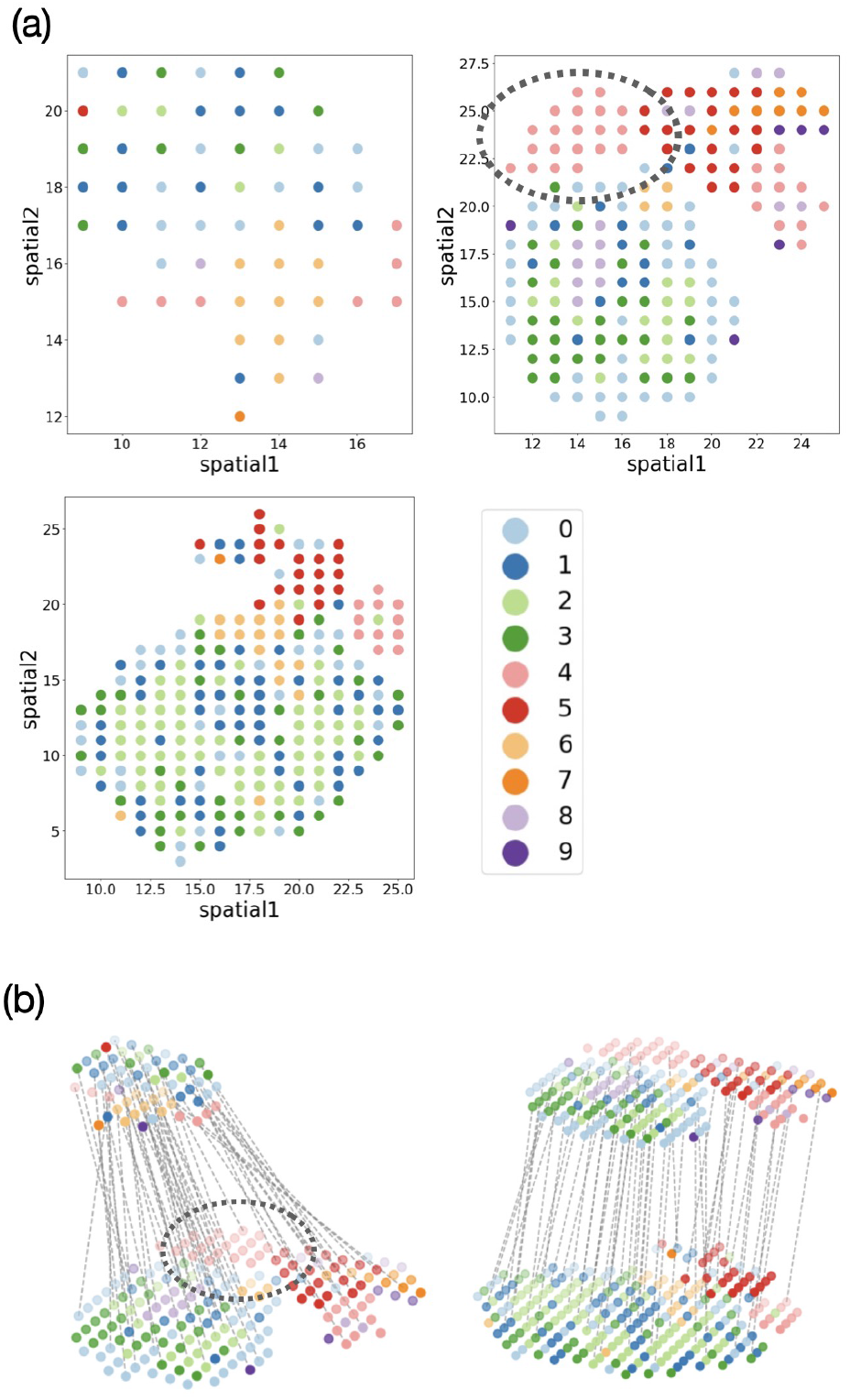
(a) Distribution of spot types in slices sampled from each developmental stage. (b) Spot matching results in ST slices using Graspot.

The distribution of spot types across spatial locations in three slices, each selected from a different developmental stage, is shown in Fig.7(b). Contrasting with recent methods that analyze this dataset by studying changes in the spatial-temporal organization of spot type composition, we employ Graspot for aligning slices across developmental time points to more intuitively understand spot relationships.

The left panel of Fig.7(c) shows the dramatic changes both on the spots number and ST slice structures. Gary circles area with almost non-existent lines means new developmental area corresponding to pink cluster 4 in Fig.7(b). The right panel of Fig.7(c) reveals that the spatial organization patterns between the two developmental stages are similar. The increase in the number of spots with age corresponds well with the heart’s growth processes.

## Discussion

In this study, we introduce Graspot, a graph attention network designed for the integration of spatial transcriptomics data using optimal transport. Graspot addresses the challenge of concurrently analyzing multiple ST slices. It generates probabilistic alignments for downstream analysis and aligns the intrinsic structures of multiple slices within a common low-dimensional space. Additionally, Graspot adeptly handles unbalanced alignment by allowing for partial overlap between slices. Our findings demonstrate Graspot’s capability to effectively perform tasks related to alignment, integration, and identification of unique spatial structures. Specifically, Graspot achieves alignments with higher accuracy compared to existing methods on ST data. Designed for scalability, Graspot can efficiently handle large-scale ST data by employing sub-network sampling and a mini-batch scheme. In this study, we have validated Graspot using 10x Genomics Visium datasets. Notably, Graspot’s application extends beyond these datasets, and it can be effectively utilized with other ST platforms such as Slide-seqV2 and Stereo-seq.

